# Molecular characterization of *Fasciola gigantica* in Punjab, Pakistan to infer the dispersal route among the neighbouring countries of the Indian subcontinent

**DOI:** 10.1101/2020.06.04.134569

**Authors:** Zia Ur Rehman, Atsushi Tashibu, Michiyo Tashiro, Imran Rashid, Qasim Ali, Osama Zahid, Kamran Ashraf, Wasim Shehzad, Umer Chaudhry, Madoka Ichikawa-Seki

**Affiliations:** Department of Parasitology, University of Veterinary and Animal Sciences, Lahore, 54200, Pakistan; Laboratory of Veterinary Parasitology, Faculty of Agriculture, Iwate University, 3-18-8 Ueda, Morioka 020-8550, Japan; Roslin Institute, University of Edinburgh, Midlothian, Scotland, EH25 9RG, United Kingdom; Department of Parasitology, Gomal University Dera Ismail Khan, Khyber Pakhtoon Khah, Pakistan

**Keywords:** Fasciola gigantica, pepck, pold, nad1, cox1

## Abstract

*Fasciola gigantica* is considered to be a major pathogen causing fasciolosis in the Indian subcontinent, resulting in millions of dollars production losses to the livestock industry. To understand the dispersal origin and the spread patterns of *F. gigantica* is important for preventing the disease. A total of 53 *Fasciola* flukes collected from buffalo and goat in the Punjab province of Pakistan, were identified as *F. gigantica* based on the multiplex PCR for the phosphoenolpyruvate carboxykinase (*pepck*) and the PCR-restriction fragment length polymorphism (RFLP) for DNA polymerase delta (*pold*). A significant genetic difference between *F. gigantica* from buffalo and goats in Pakistan was indicated by the genetic analysis of two distinct mitochondrial markers [NADH dehydrogenase subunit 1 (*nad1*) and cytochrome C oxidase subunit 1 (*cox1*)]. Phylogenetic analysis of the seventeen *nad1* haplotypes of *F. gigantica* from Pakistan with those in neighbouring countries of the Indian subcontinent revealed that all the haplotypes were clustered in haplogroup A. *Fasciola gigantica* with the eight haplotypes might be expanded in Pakistan from Indian origin, along with the migration of the domestic animals, since they were related to Indian haplotypes. In contrast, the remaining nine haplotypes were not shared with any neighbouring countries, suggesting independent origin, or possibly come from neighbouring Middle East countries. Our study provides a proof of concept for a method that could be used to investigate the epidemiology of *F. gigantica* regarding the development of sustainable parasite control strategies.

## 1. Introduction

Fasciolosis, caused by the liver fluke of genus *Fasciola*, is a neglected zoonotic disease that results in a severe economic losses in the livestock industry (Aghayan et al., 2019). Fasciolosis is one of the most widely spread diseases reported from over 50 countries mostly in Asia, Africa and America (Mas-Coma, 2003; Mas-Coma et al., 2005; Toledo and Fried, 2014). The incidents of fasciolosis have increased over the past two decades, possibly because of the changes in farming practices, development of anthelmintic resistance and climatic changes (Sabourin et al., 2018). The genus *Fasciola* comprises of two important species*. Fasciola hepatica* is found in temperate zones, whereas *F. gigantica* is generally considered to be a parasite of tropical areas (Mas-Coma et al., 2009). Both species co-exist in subtropics, which can result in the formation of intermediate or hybrid forms mainly in Asian countries (Ichikawa and Itagaki, 2010b; Rokni et al., 2010). *Fasciola* infects the livers and bile ducts of ruminants and other mammals, while the snails of Lymnaeidae family act as their intermediate hosts (Toledo and Fried, 2014; Usip et al., 2014).

Mitochondrial markers have been used for the phylogenetic characterizations of *Fasciola* species to examine the propagation route of this group of parasites in many countries (Ai et al., 2011; Elliott et al., 2014; Ichikawa-Seki et al., 2017; Ichikawa and Itagaki, 2012; Semyenova et al., 2006; Thang et al., 2019). In the Asian subcontinent, *F. gigantica* has been divided into three haplogroups. Haplogroups B and C have been predominant mainly in Southeast Asian countries including Thailand (Chaichanasak et al., 2012), Myanmar (Ichikawa et al., 2011) and Indonesia (Hayashi et al., 2016). Haplogroup A has been distributed in the Indian subcontinent including East India (Hayashi et al., 2015), Bangladesh (Mohanta et al., 2014), Nepal (Shoriki et al., 2014) and Myanmar (Ichikawa et al., 2011). The zebu cattle (*Bos indicus*) and water buffalo (*Bubalus bubalis*) have been considered to be the definitive host of *F. gigantica* (Kikkawa et al., 2003). Hayashi et al. (2015) demonstrated that the haplogroup A of *F. gigantica* originated in the Indus River vally, and the unregulated animal movement might be involved in the spread of this parasite species. However, limited information is available on *F. gigantica* fluke in Pakistan to reveal the expansion history of this parasite through Indus River vally of the Indian subcontinent.

In this paper, we describe a study using liver flukes collected from buffalo and goat slaughtered in abattoirs in the Punjab province of Pakistan, with the following aims: i) to perform the species identification using the most reliable nuclear markers: phosphoenolpyruvate carboxykinase (*pepck*) and DNA polymerase delta (*pold*) genes (Shoriki et al., 2016); ii) to determine the propagation route of *F. gigantica* in the Indian subcontinent using mitochondrial NADH dehydrogenase subunit 1 (*nad1*) and cytochrome C oxidase subunit 1 (*cox1*) genes.

## 2. Materials and Methods

### 2.1. Sample collection and gDNA extraction

A total of 14 *Fasciola* infected livers (7 buffalo and 7 goats) were collected from the animals slaughtered at the Punjab Agriculture & Meat Company (PAMCO) Lahore (31.4330° N, 74.1945° E). The livers were transported from abattoir to the laboratory on ice, where flukes were recovered from the biliary ducts by dissection. A total of 53 flukes (25 from 7 buffalo and 28 from 7 goats) (Table 1) were thoroughly washed with phosphate-buffered saline (PBS), and preserved in 70% ethanol until use. A small portion of the vitelline glands from the posterior part of each fluke was used for DNA extraction using the High Pure DNA Extraction Kit (Roche, Mannheim, Germany) following the manufacturer’s protocols, and stored at −20°C until further use.

**Table 1:**
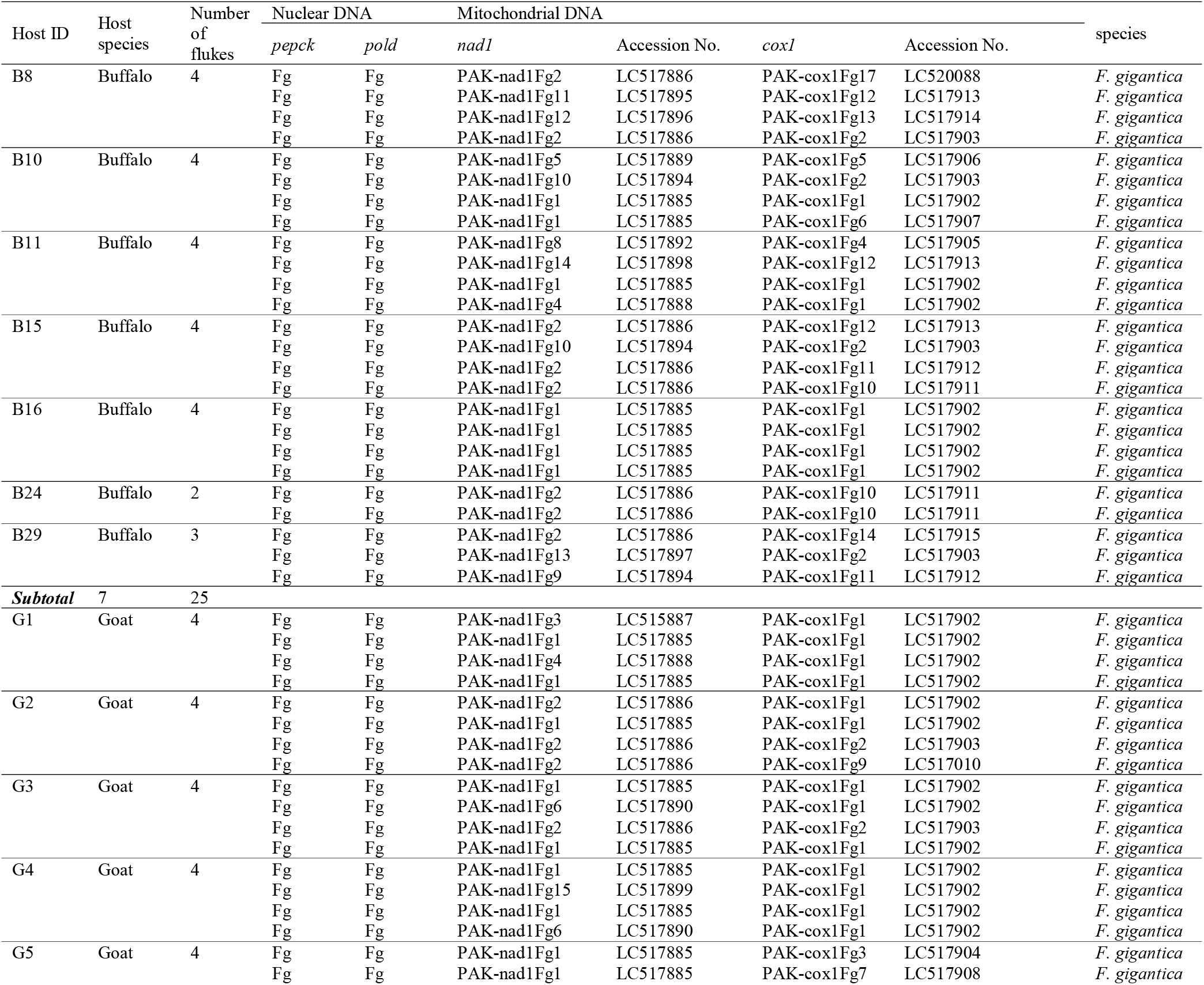
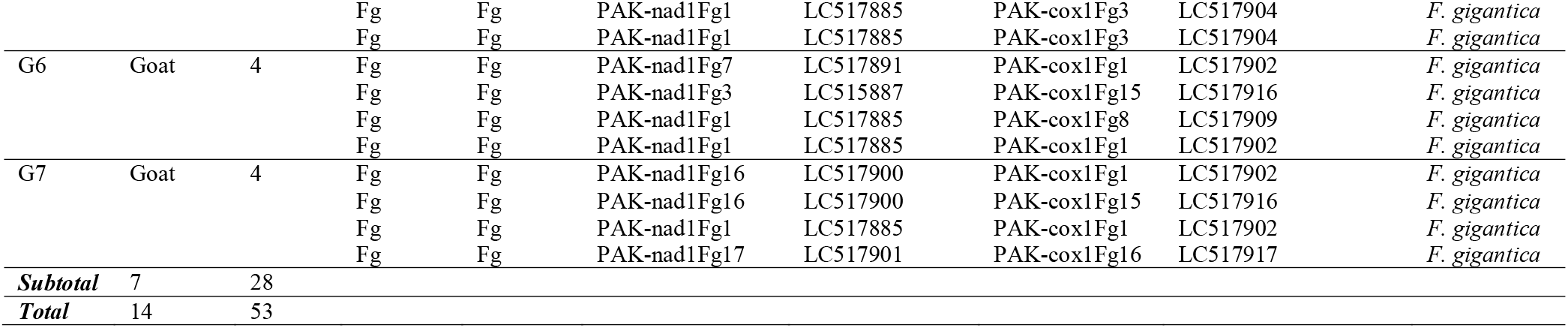
Profiles of *Fasciola* flukes used in this study.

### 2.2. *Multiplex PCR and PCR-RFLP of pepck and pold genes of* Fasciola *species*

The fragments of *pepck* were amplified through the multiplex PCR assay previously described by Shoriki et al. (2016). The PCR amplicons were run on 1.8% agarose gels for 30 min to detect the *F. hepatica* (approximately 500bp), *F. gigantica* (approximately 240bp) or hybrid fragment patterns (both the fragments). The fragments of *pold* were analysed through the PCR-RFLP assay previously described by Shoriki et al. (2016). The PCR products were subsequently digested with *Alu*I enzyme (Toyobo, Osaka, Japan) at 37 °C for three hours. The resultant products were run on 1.8% agarose gels for 30 min to detect the *F. hepatica* (approximately 700bp), *F. gigantica* (approximately 500bp) or hybrid fragment patterns (both the fragments).

### 2.3. PCR amplification and sequencing of nad1 and cox1 genes

The fragments of *nad1* (535bp) and *cox1* (430bp) genes were amplified through a PCR assay previously described by Itagaki et al. (2005). The PCR amplicons were purified using NucleoSpin Gel and PCR Clean-up Kit (Macherey-Nagel, Düren, Germany) and sequenced by Eurofin Genomics K.K. (Tokyo, Japan). The resultant DNA sequences were assembled using ATGC v. 6.0.3 (Genetyx Co., Tokyo, Japan) and the haplotypes were identified by GENETYX v. 10.0.2 (Genetyx Co.).

### 2.3. Median-joining network and diversity indices of nad1 and cox1 genes

Median-joining (MJ) network was constructed using Network v. 5.0.1.1 software (Tajima, 1989) to determine the phylogenetic relationships among the *nad1* and *cox1* haplotypes. Median-joining (MJ) network has been further used to compare the *nad1* haplotypes of the present study with the reference *nad1* haplotypes of *F. gigantica* from India (Ichikawa and Itagaki, 2010a), Bangladesh (Mohanta et al., 2014), Nepal (Shoriki et al., 2014), Myanmar (Ichikawa et al., 2011), Thailand (Chaichanasak et al., 2012), Vietnam (Itagaki et al., 2009), Indonesia (Hayashi et al., 2016), China (Peng et al., 2009), Korea (Ichikawa and Itagaki, 2012) and Japan (Itagaki et al., 2005). The reference *nad1* haplotypes were retrieved from the GenBank and their frequencies were referred from the previous reports.

The diversity indices, including the number of variable sites (*S*), the number of haplotypes (*h*), and the nucleotide diversity (*π*) were calculated using DnaSP software v. 5.1 (Librado and Rozas, 2009). Tukey’s test was performed by GraphPad Prism v. 7.04 (GraphPad Software Inc., San Diego, CA, USA) to detect the significant differences in *π* values among the populations. First, the samples of the present study were compared to each other to find the differences between buffalo and goat. In the next step, the indices of *nad1* were compared with those of the reference populations of *F. gigantica* from India (Hayashi et al., 2015), Bangladesh (Mohanta et al., 2014), Nepal (Shoriki et al., 2014) and Myanmar (Ichikawa et al., 2011) to find relationships between the neighbouring countries.

The pairwise fixation index (*F*_ST_) values between the *F. gigantica* samples derived from buffalo and goat were calculated using Arlequin program v. 3.5.2.2 (Loftus et al., 1994) to find genetic differences. If the *F*_ST_ values approaching 1 indicate extreme genetic differentiation between the two populations.

## Results

### 3.1. Species identification

The fragment analysis by the multiplex PCR and the PCR-RFLP for *pepck* and *pold* showed that a total of 53 flukes collected from buffalo and goat were *F. gigantica* (Table 1). These results are complemented with the previous reports which suggest that *F. gigantica* is the predominant species in Punjab province of Pakistan (Chaudhry et al., 2016; Rehman et al., 2020).

### 3.2. Mitochondrial haplotype distribution of F. gigantica between buffalo and goat

The haplotype distribution was analysed separately for 53 individual *F. gigantica* flukes. A total of seventeen *nad1* haplotypes were detected in *F. gigantica* (Table 1). The MJ network revealed that PAK-nad1Fg1 and PAK-nad1Fg2 were the two predominant haplotypes present in both buffalo and goat (Fig. 1A) with four nucleotide substitutions between them. A total of seventeen *cox1* haplotypes were detected in *F. gigantica* (Table 1). The MJ network revealed that PAK-cox1Fg1 and PAK-cox1Fg2 were the predominant haplotypes found in both buffalo and goat (Fig. 1B), with three nucleotide substitutions between them.

**Fig. 1:**
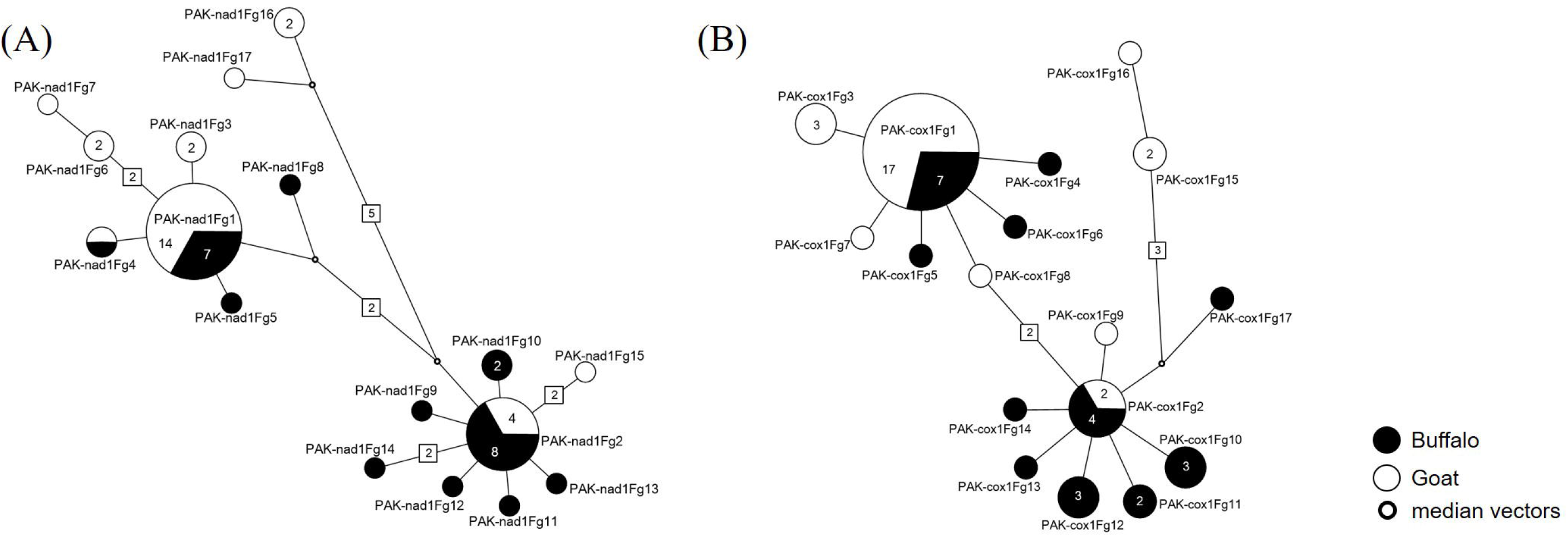
Median-joining network based on the mitochondrial (A) *nad1* and (B) *cox1* haplotypes of *F. gigantica* from Pakistani origin. Each circle indicates a single haplotype. Black colour indicates the haplotypes from Buffalo and white colour haplotypes from the goat. Small circles are the median vectors needed to connect the haplotypes.

The *π* for *nad1* and *cox1* were compared between *F. gigantica* samples acquired from both hosts. For the *nad1* gene, a higher *π* value was observed in the *F. gigantica* populations obtained from goat, but the result was opposite for the *cox1* gene (Table 2), and therefore, more diverse i.e. older population could not be determined. The *F*_ST_ values between the two hosts for both the genes *(nad1:* 0.12931, *cox1:* 0.16914) were statistically significant (*P* < 0.05), indicating the existence of genetic differentiation.

**Table 2:** Diversity indices of *F. gigantica* populations based on the nucleotide sequences of mitochondrial markers.

| Gene | Host | *N* | *S* | *h* | $\pi$ | *SD* |
| --- | --- | --- | --- | --- | --- | --- |
| *nad1* | Buffalo | 25 | 14 | 11 | 0.00533 | 0.00049 |
|  | Goat | 28 | 17 | 9 | 0.00668 | 0.00135 |
| *cox1* | Buffalo | 25 | 12 | 11 | 0.00612 | 0.00047 |
|  | Goat | 28 | 10 | 8 | 0.00493 | 0.00125 |

*N*: number of flukes accessed; *S*: number of variable sites; *h*: number of haplotypes; $\pi$ : nucleotide diversity. All values were statistically significant ( $P < 0.05$ ).

### 3.3. Comparative analysis of F. gigantica with that from neighboring countries

The MJ network analysis of seventeen *nad1* haplotypes revealed that PAK-nad1Fg2 haplotype had an identical sequence with the *F. gigantica* haplotypes from India (ND1-E6), Nepal (Fg-ND1-N1), Myanmar (Fg-M15), Bangladesh (Fg-NDI-Bd9) and Thailand (Fg-ND1-Thai13) (Fig. 2). The PAK-nad1Fg9, 10, 11, 12, 14 and 15 haplotypes had a single or double nucleotide substitutions present nearly to the PAK-nad1Fg2 haplotype. The PAK-nad1Fg13 halotype had an identical sequence with the *F. gigantica* haplotypes from India (ND1-E7) and Myanmar (Fg-M16), and PAK-nad1Fg10 was identical to ND1-IN14 haplotype from India (Fig. 2). In contrast, the remaining nine haplotypes including PAK-nad1Fg1 (a most dominanted haplotype) were not related to any reference haplotypes from neighbouring countries (Fig. 2).

**Fig. 2:**
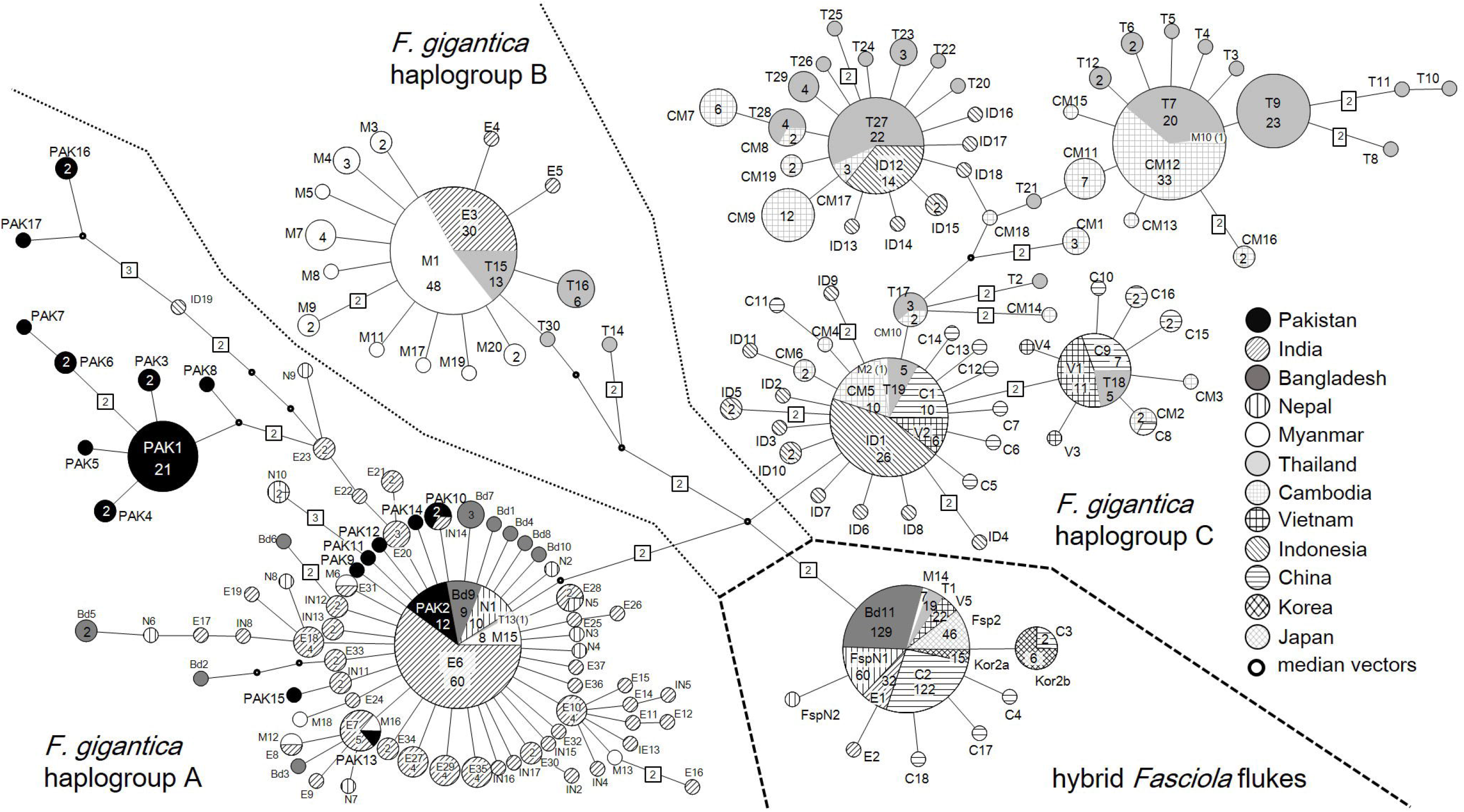
Median-joining network based on the mitochondrial *nad1* haplotypes of *F. gigantica* in Pakistan and other countries. The *Fasciola* flukes from Pakistan are shown in black colour. Each circle indicates a single haplotype. Small circles are the median vectors which are needed to connect the haplotypes. The haplotype codes are shown within or adjacent to the circles. Numbers on each circle and node indicate the number of flukes and the number of substitution sites, respectively.

The *π* values for *nad1* were compared between *F. gigantica* samples from Pakistan and neighbouring countries. The data suggest that the *F. gigantica* of the present study had the highest *π* value among the populations (Table 3), implying a higher nucleotide diversity in Pakistani populations as compared to the neighbouring countries.

**Table 3.**
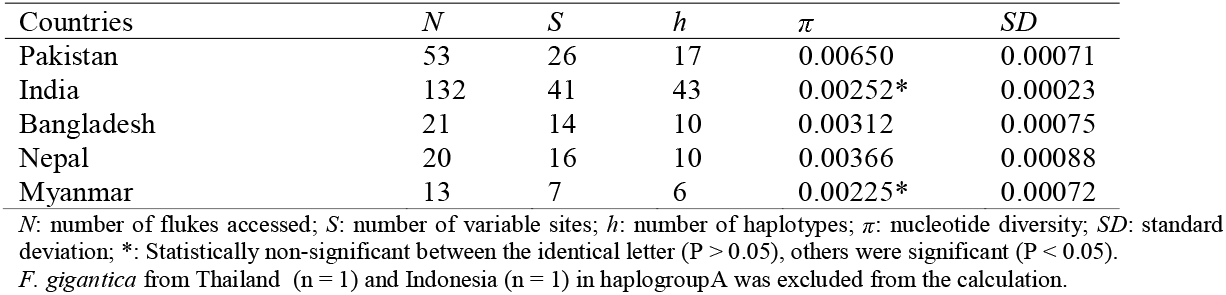
Diversity indices of *F. gigantica* within haplogroup A based on the nucleotide sequences of nad1 gene.

## Discussion

Historically, it has been suggested that *F. gigantica* might originate and spread by zebu cattle (*Bos indicus)* and water buffalo (*Bubalus bubalis*) in the Indian subcontinent (Peng et al., 2009). The water buffalo was domesticated in the Indus River valley (modern-day Pakistan) at around 7,000 to 8,000 BC, whereas zebu cattle was domesticated in the Indus Valley, Near and Middle East and Eastren Europe around 5,000 years ago (Bradley et al., 1996; Loftus et al., 1994; Tanaka et al., 1996). Since Pakistan is a part of the Indian subcontinent, free movement of zebu cattle and water buffalo probably play a significant role in the spread of *F. gigantica* in the region. Over the past few decades, high levels of animal movement have been reported in domestic ruminants in the Indian subcontinent (Kelley et al., 2016; Vilas et al., 2012). The farmers rear multiple species of animals to meet their livelihood in this region (Devendra, 2007). The mixed farming system might play an important role in the spread of *F. gigantica*. The animal movement patterns differ between farms, and *F. gigantica* infects a wide range of hosts including domestic and wild animals. Therefore, human activities potentially enable the spread of this parasite (Rojo-Vázquez et al., 2012). Hence genetic analysis are needed to understand the corresponding origin and spread of *F. gigantica* infections, which aid in the development of parasite control strategies (Hayashi et al., 2016).

In the present study, a significant genetic differentiation between *F. gigantica* samples from buffalo and goat was suggested by the *F*_ST_ value, which might be due to the difference in host immunity (Haroun and Hillyer, 1986; Piedrafita et al., 2004; Roberts et al., 1997) or due to the variances in the geographical position of these two hosts in the Punjab province of Pakistan. Generally, the lowlands of Punjab are more prone to flooding and reported to be highly populated with buffalo, in comparison, goat are resided in higher areas or keep moving to different areas due to human travelling (Afshan et al., 2014). However, the expansion history of *F. gigantica* between the hosts could not be inferred in the present study; the older host in Punjab could not be determined since the opposite statistical differences were observed in the *π* values of *nad1* and *cox1*.

The current study revealed that at least two origins of *F. gigantica* in Pakistan with reference to the neighbouring countries. In haplogroup A, the eight *F. gigantica* haplotypes of the present study had a close relationship with the haplotypes from India, Bangladesh, Nepal, Myanmar and Thailand. The MJ network indicates that the haplotypes detected in India were apparently more diverse than Pakistan, which indicates the hypothesis of the expansion of these haplotypes from India to Pakistan. In contrast, the nine Pakistani haplotypes were not shared with any neighbouring countries, suggesting an independent origin, or possibly come from neighbouring Middle East countries where genetic analysis of *F. gigantica* has never been conducted. Further studies from different areas of Pakistan, as well as neighbouring Middle East countries, will be required to reveal the origin and dispersal direction of these haplotypes.

In summary, the present study provides preliminary insights into the origin and spread of *F. gigantica* in Pakistan and the neighbouring countries of the Indian subcontinent. We have also described the genetic difference between *F. gigantica* populations derived from buffalo and goat. Overall, the study provides a benchmark and opens a new avenue for more detailed analysis in this region. It might be helpful to involve higher samples size, more host species from different areas to get more conclusive results of the genetic diversity of *F. gigantica* among buffalo, goat, sheep and cattle as well as their possible spread patterns in the country and the subcontinent.

## Acknowledgements

The authors acknowledge Mr Zain Ali khan, Mr Obaid Mohammad Abdullah and Mr Muhammad Omer Gulzar for their support and participation during sample collection. This work was supported by a Grant-in-Aid for Scientific Research (B) from MEXT KAKENHI (Grant Number16H05806).

